# MFF SUMOylation regulates neuronal mitochondrial morphology and facilitates AMPK-induced mitochondrial fragmentation

**DOI:** 10.64898/2026.09.07.749864

**Authors:** Richard Seager, Daniela Franchini, Jeremy M. Henley, Kevin A. Wilkinson

## Abstract

Mitochondrial dysfunction and fragmentation in neurons are characteristics of many neurodegenerative conditions. AMP-activated protein kinase (AMPK) is a cellular energy sensor which is hyperactivated in Alzheimer’s disease. Under stress conditions, AMPK-mediates phosphorylation of the outer mitochondrial membrane protein mitochondrial fission factor (MFF), leading to SUMOylation at Lys^151^. MFF SUMOylation promotes fission and subsequent mitochondrial fragmentation. Here, using primary neuronal culture, we investigate the role of MFF SUMOylation in mitochondrial morphology and function. We show that MFF SUMOylation is required to maintain neuronal mitochondrial length under basal conditions in axons and dendrites. Using the AMPK activator (AICAR) to induce mitochondrial fission, show that preventing MFF SUMOylation reduces AICAR-induced fragmentation of dendritic mitochondria. These findings reconcile observations of AMPK hyperactivation and mitochondrial fragmentation by revealing the AMPK-MFF-SUMO pathway to be crucial in neuronal mitochondrial fission.

## Introduction

Neurons are anatomically complex, energy-demanding cells, that contain distinct compartments with specific functions and biochemical requirements. Synapses in particular are energy intensive sites, and mitochondria are crucial for providing the ATP required to power synaptic transmission (Devine & Kittler, 2018; Tong *et al*, 2026) and to buffer calcium which modulates neurotransmission (Walters & Usachev, 2023; Datta & Jaiswal, 2021). Mitochondria in neurons constantly move, fuse and divide (Cagalinec *et al*, 2013; Chang *et al*, 2006), termed mitochondrial dynamics, to monitor and maintain the correct morphology, localisation and quality control, to ensure a healthy mitochondrial population (Seager *et al*, 2020).

Neuronal activity increases mitochondrial localisation at both pre-and postsynaptic synaptic sites (Li *et al*, 2004; Vaccaro *et al*, 2017; Obashi & Okabe, 2013; Chang *et al*, 2006). This enhanced synaptic positioning of mitochondria is key to integrate mitochondrial metabolism with neuronal activity. At the pre-synapse, local mitochondria regulate synaptic vesicle release by modulating calcium levels (Vaccaro *et al*, 2017) and providing sufficient ATP during activity (Sun *et al*, 2013; Verstreken *et al*, 2005). At the post-synapse, local mitochondria are critical for powering protein synthesis and spine plasticity (Bapat *et al*, 2024; Rangaraju *et al*, 2019).

Moreover, neuronal activity underpins changes in mitochondrial dynamics. Decreased neuronal activity causes dendritic mitochondria to elongate (Hatsuda *et al*, 2023; Chang *et al*, 2006; Virga *et al*, 2024), whereas increased activity induces fission (Li *et al*, 2004; Divakaruni *et al*, 2018). Interestingly, axonal mitochondria appear to be less morphologically responsive to changes in neuronal activity (Hatsuda *et al*, 2023; Chang *et al*, 2006), although axonal mitochondrial shortening has been observed following increased neuronal activity (Chang *et al*, 2006).

Reciprocally, mitochondrial health and size have profound effects on neuronal function and spine activity. Fragmentation of dendritic mitochondria blocks newly synthesised proteins (Rangaraju *et al*, 2019) and fails to support spine growth in stimulated spines (Bapat *et al*, 2024). Elongated axonal mitochondria sequester more calcium, resulting in reduced neurotransmitter release and decreased axonal branching (Lewis *et al*, 2018). The inextricable link between mitochondrial dynamics and neuronal function is underscored by multiple neurodegenerative diseases associated with dysfunctional mitochondrial dynamics (Chen *et al*, 2023; Sirois *et al*, 2026).

The AMP-activated protein kinase (AMPK) is the major energy sensor of the cell, which becomes activated when the ATP/AMP ratio decreases. Activated AMPK restores energy homeostasis by promoting energy-generating processes and inhibiting energy-demanding processes (Herzig & Shaw, 2018). The outer mitochondrial membrane protein mitochondrial fission factor (MFF) is the primary receptor for the cytosolic protein dynamin-related protein 1 (DRP1). Upon recruitment to MFF, DRP1 forms helices around the mitochondria to facilitate mitochondrial fission (Otera *et al*, 2016; Losón *et al*, 2013). Following severe bioenergetic stress, AMPK phosphorylates MFF at Ser^155^ and Ser^172^ (Ducommun *et al*, 2015; Toyama *et al*, 2016), which is necessary and sufficient to promote fragmentation (Toyama *et al*, 2016). We previously identified MFF as a target of the small ubiquitin-like modifier (SUMO) at Lys^151^, and showed that following phosphorylation, MFF SUMOylation is enhanced, which promotes stress-induced mitochondrial fission in MEF cells (Seager *et al*, 2024).

Neuronal AMPK is rapidly activated during synaptic activity, which upregulates glycolysis and mitochondrial respiration to meet the energy requirements for synaptic plasticity and learning and memory (Marinangeli *et al*, 2018), and is required for mitochondrial recruitment to pre-synaptic sites to sustain ATP levels during neuronal activity (Li *et al*, 2020). However, chronic AMPK activation leads to loss of synaptic proteins in SH-SY5Y cell lines (Yang *et al*, 2022), primary neurons and *in vivo*, corresponding to a functional decrease in synaptic activity (Domise *et al*, 2019). Moreover, the AMPK-induced mitochondrial fragmentation has been reported as a prerequisite for spine loss (Lee *et al*, 2022), suggesting that maintaining mitochondrial integrity and preventing fission blocks mitophagic clearance and subsequent spine loss. These findings indicate that regulated AMPK activity is crucial for long-term synaptic function, and prolonged AMPK activity is detrimental to synaptic function and integrity.

AMPK is hyperactivated in Alzheimer’s disease (AD) (Vingtdeux *et al*, 2011; Ma *et al*, 2014; Fang *et al*, 2019). Furthermore, the toxic protein amyloid-β (Aβ), implicated in AD pathology, activates AMPK in primary neurons, leading to mitochondrial fragmentation, mitophagy, and spine loss (Lee *et al*, 2022; Thornton *et al*, 2011; Mairet-Coello *et al*, 2013; Domise *et al*, 2019). Excessive mitochondrial fragmentation and mitochondrial dysfunction are hallmarks in AD pathology (Fang *et al*, 2019; Wang *et al*, 2016, 2008). In a variety of *in vitro* and *in vivo* neurodegenerative models, reducing mitochondrial fragmentation is neuroprotective (Strucinska *et al*, 2025; Lee *et al*, 2022; Filichia *et al*, 2016; Bido *et al*, 2017; Qi *et al*, 2012; Grohm *et al*, 2012), indicating that defining ways to modulate mitochondrial dynamics could be a powerful strategy for neuroprotection.

In this study, we expressed a series of MFF phosphorylation and SUMOylation mutants in primary rat neurons. We show that MFF SUMOylation is a key modification required for regulating neuronal mitochondrial morphology under basal conditions. AMPK activation with AICAR induces mitochondrial fragmentation in dendrites but not in the axons. Critically, SUMOylation-deficient MFF-expressing neurons are resistant to AICAR-induced mitochondrial fission in dendrites. Together, these results demonstrate that MFF-SUMOylation is key for regulating basal mitochondrial size and AMPK-mediated mitochondrial remodelling in dendrites.

## Results

### AMP-mimetic AICAR fragments dendritic mitochondria and reduces mitochondrial function

To model the chronic AMPK activation observed in AD, we treated primary neurons with the AMP-mimetic AICAR to activate AMPK. We transfected primary hippocampal neurons with mitoDS-red at DIV (days in vitro) 15 to label the mitochondria to measure mitochondrial morphology changes under AICAR conditions. Three days post-transfection, neurons were treated with 1 mM AICAR for three days (DIV 18-21), as used previously (Domise *et al*, 2019). Cells were then fixed and stained for ankyrin-G (marker of the axonal initial segment), to distinguish between axonal and dendritic processes (Fig 1A). In control cells, the axon exhibited short, sparsely distributed mitochondria, compared to elongated mitochondria in the dendritic processes, as previously reported (Lewis *et al*, 2018; Chang *et al*, 2006). Following AICAR treatment, we observe a significant 16.9% decrease in average dendritic mitochondrial length (Fig 1B), and a corresponding shift in the distribution of dendritic mitochondrial lengths towards shorter mitochondria (Fig 1C). Interestingly, no effect was observed in the mean mitochondrial length in the axon or in their distribution, suggesting a compartment-specific effect of AMPK activation (Fig 1B, C). To confirm AICAR activates AMPK in our neuronal culture system, we treated primary cortical neurons with 1 mM AICAR at DIV 18 for three days. Total protein lysates were then immunoblotted for AMPK activation as assessed by AMPK phosphorylation (Fig 1D), which validated AMPK activation by AICAR.

**Figure 1.**
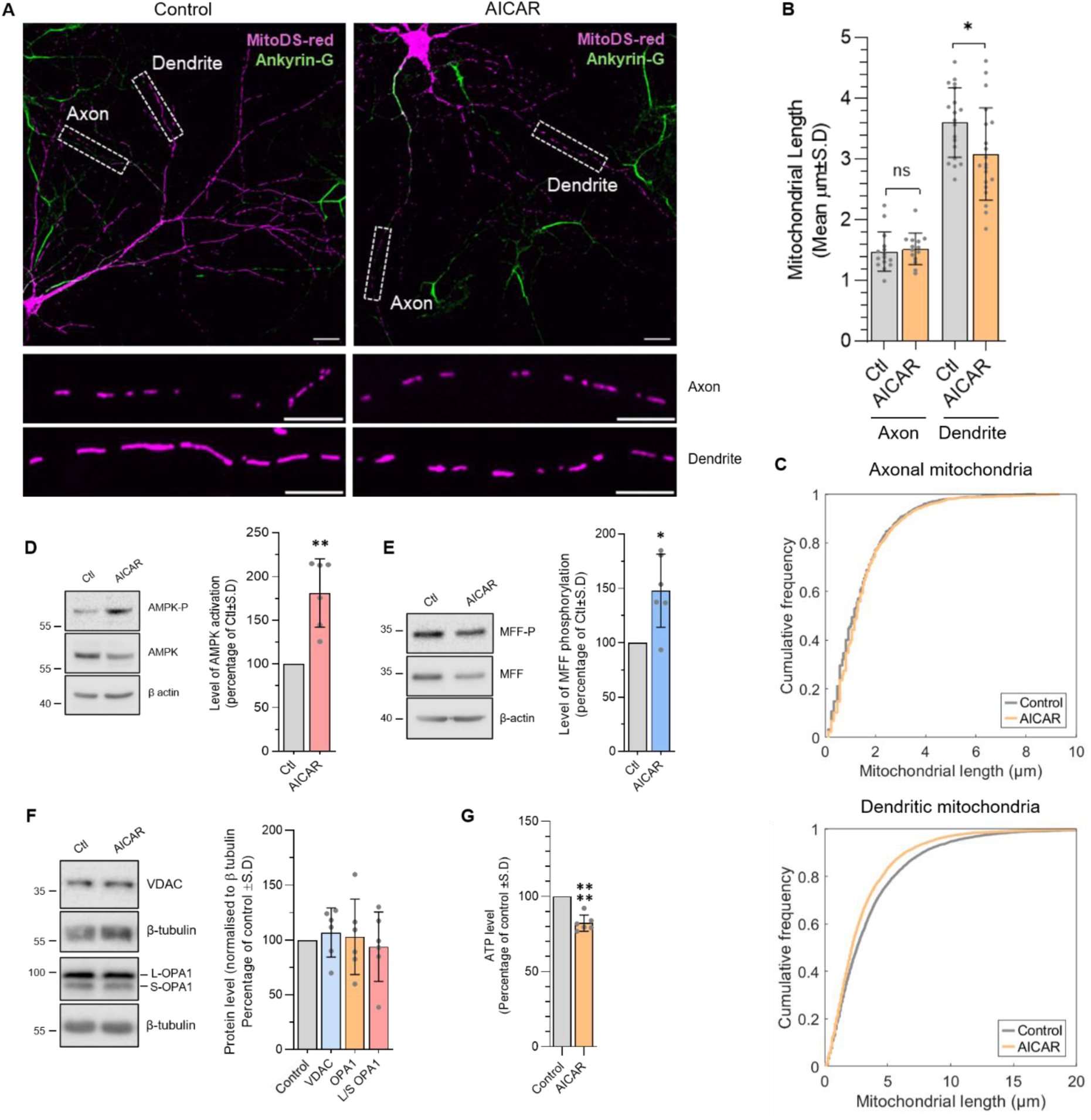
AICAR fragments mitochondria specifically in the dendrites. (**A**) Confocal images of primary hippocampal neurons transfected with mito-DSred and treated with 1 mM AICAR for 72hrs (DIV 18-21). Neurons were fixed and stained for ankyrin-G to define the axonal initial segment. Scale bar 20µm, enlargements show highlighted region of axon and dendrite, scale bar 10µm. (**B**) Quantification of mean mitochondrial lengths. Data gathered from three independent experiments. Two sample t test (dendrites), p*<0.05, Mann-Whitney U test (axons) showed no significance. Cumulative frequency of all axonal and dendritic mitochondria shown in **C**. (**D**) AICAR-treated (1 mM, 72h) cortical neuronal protein lysate was probed for phosphorylated AMPK and total AMPK levels. Quantification shows AMPK activation (AMPK-P/AMPK), expressed as percentage of control. Data gathered from 6 independent experiments, one sample t-test, p**<0.01. (**E**) AICAR-treated (1 mM, 72h) cortical neurons were probed for MFF at S172 and total MFF. Quantification shows level of MFF phosphorylation (MFF-P/MFF), expressed as percentage of control. Data gathered from 6 independent experiments, one sample t-test, p*<0.05. (**F**) AICAR-treated cortical neurons were probed for VDAC and OPA1. Quantification shows data normalised to β-tubulin or the ratio of lower band/higher band (short/long OPA1) and expressed as percentage of control. Data generated from 6 independent experiments. One sample t test used, no significance difference detected. (**G**) Mitochondrial function assay. Primary cortical neurons ATP levels were measured following AICAR treatment (1mM, DIV18-21). Data expressed as percentage of control, 6 independent experiments, one sample t-test, p****<0.0001.

AMPK phosphorylates MFF to promote fission (Toyama *et al*, 2016) so we probed for phosphorylated MFF, which was significantly increased by almost 50% when normalised to total MFF levels (Fig 1E). To examine whether there is a loss of mitochondrial mass, we assessed the expression of the mitochondrial marker VDAC, which remained unchanged by AICAR treatment (Fig 1F). Moreover, there was no change in total levels of the fusion protein OPA1, or in the ratio of long to short OPA1 isoforms (increase in short OPA1 is associated with fission (Anand *et al*, 2014)) (Fig 1F). These data suggest the observed fragmentation is an up-regulation of fission, rather than reduced fusion. Finally, we probed for the GTPase DRP1, which drives mitochondrial fission and is the ligand of MFF. Levels of both DRP1 and MFF were reduced by AICAR (Fig S1A). These results are surprising, since the mitochondria are undergoing fission, and we postulate these changes to be a compensatory mechanism to prevent excessive fragmentation.

Since mitochondrial morphology and function are inextricably linked, we next assessed the effect of AICAR on mitochondrial function in cortical neurons by measuring ATP levels. Neurons treated with AICAR exhibited a significant ∼20% reduction in ATP levels compared to untreated cells (Fig 1G). These neurons were cultured in standard high glucose neurobasal media (25 mM), so they likely primarily generated ATP using glycolysis rather than mitochondrial oxidative phosphorylation, and therefore the mitochondria may be less responsive. To promote aerobic respiration, we cultured neurons in low glucose (5 mM). The AICAR challenge in the low glucose cells elicited a similar decrease in ATP levels (Fig S1B), indicating that the decrease observed is due to a decrease in mitochondrially generated ATP, even in the presence of high glucose. Together, these results indicate that neuronal AMPK activation phosphorylates MFF, which induces dendritic mitochondrial fragmentation, but not in the axon, and loss of function.

### MFF SUMOylation regulates basal neuronal mitochondrial length

MFF SUMOylation is enhanced following mitochondrial stress, AMPK activation and AICAR treatment (Seager *et al*, 2024). Building on this observation of AICAR-mediated neuronal mitochondrial remodelling, we tested the possible role of MFF SUMOylation. We transfected primary hippocampal neurons with mito-DSred together with an empty CFP plasmid, wild-type CFP-MFF or MFF SUMO-deficient mutant K151R CFP-MFF (Fig 2A). We also included a mito-DSred alone condition as a control to confirm overexpression of CFP does not induce any adverse effects.

**Figure 2.**
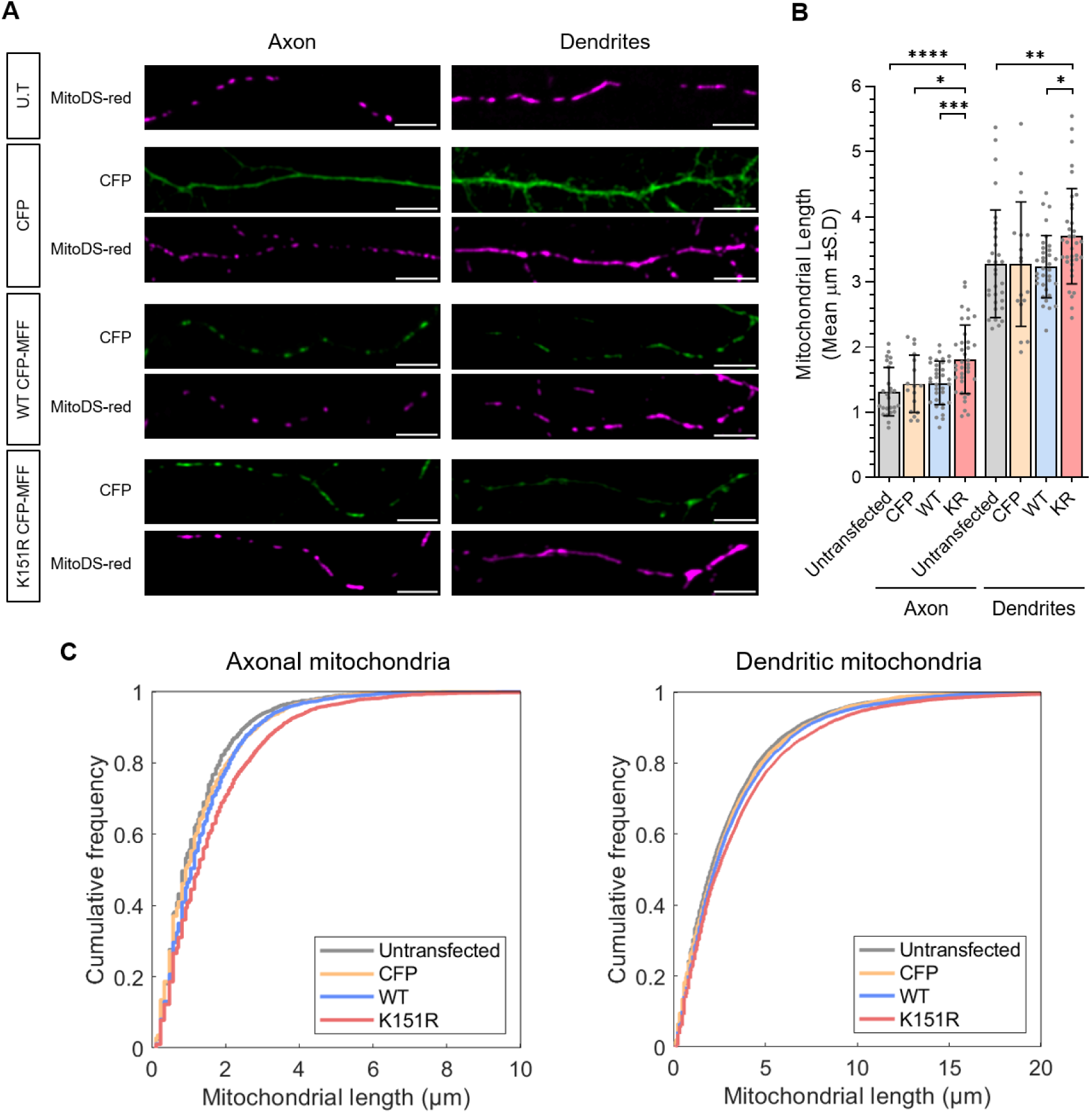
MFF SUMOylation regulates neuronal mitochondrial size. (**A**) Confocal images of primary hippocampal neurons transfected with mito-DSred along with the indicated CFP-MFF constructs, or mito-DSred alone (termed untransfected, U.T). Neurons were fixed and stained for GFP and ankyrin-G (to define the axonal initial segment). Images show mitochondria from the axon and dendrites, scale bar 10µm. (**B**) Quantification of mean mitochondrial lengths per cell. Data generated from 3-5 independent experiments. One-Way ANOVA followed by Tukey’s multiple comparison test used to determine significance, p*<0.05, p**<0.01, p***<0.005, p****<0.0001. (**C**) Cumulative frequency of all axonal and dendritic mitochondria.

There was no difference in the mean mitochondrial length in axons and dendrites between mito-DSred alone, CFP or wild-type MFF. There was, however, a significant elongation of average axonal mitochondria (of 24.9%, 26.2% and 37.7%, when compared to mito-DSred alone, CFP or wild-type MFF, respectively) on expression of non-SUMOylatable K151R MFF (Fig 2B). Likewise, the non-SUMOylatable K151R MFF elicited a significant increase in dendritic mitochondrial length compared to mitoDS-red and compared to wild-type MFF expressing cells (∼14.4% increase, Fig 2B).

Taking all mitochondria measured and expressing as a cumulative frequency, there was a shift in the distribution towards longer mitochondria in cells expressing K151R, in both the axonal and dendritic compartments (Fig 2C). Together, these results demonstrate that MFF SUMOylation basally regulates both axonal and dendritic mitochondrial length, and that blocking MFF SUMOylation enhances mitochondrial length.

### MFF phosphorylation-SUMOylation axis regulates basal neuronal mitochondrial length

To further investigate the sequential phosphorylation-SUMOylation modification axis of MFF on neuronal mitochondrial morphology, we co-expressed mito-DSred with the following CFP-MFF constructs: wild-type, K151R, the AMPK phosphorylation site mutants 2SA: double phospho-null, and 2SD: double phospho-mimetic, and KR/2SD: SUMO mutant (K151R) plus the double phospho-mimetic mutation (Fig 3A). The sites of the serine substitution to alanine (A) or aspartic acid (D) are based on the identified AMPK sites of MFF Ser^155^ and Ser^172^ (Toyama *et al*, 2016).

**Figure 3.**
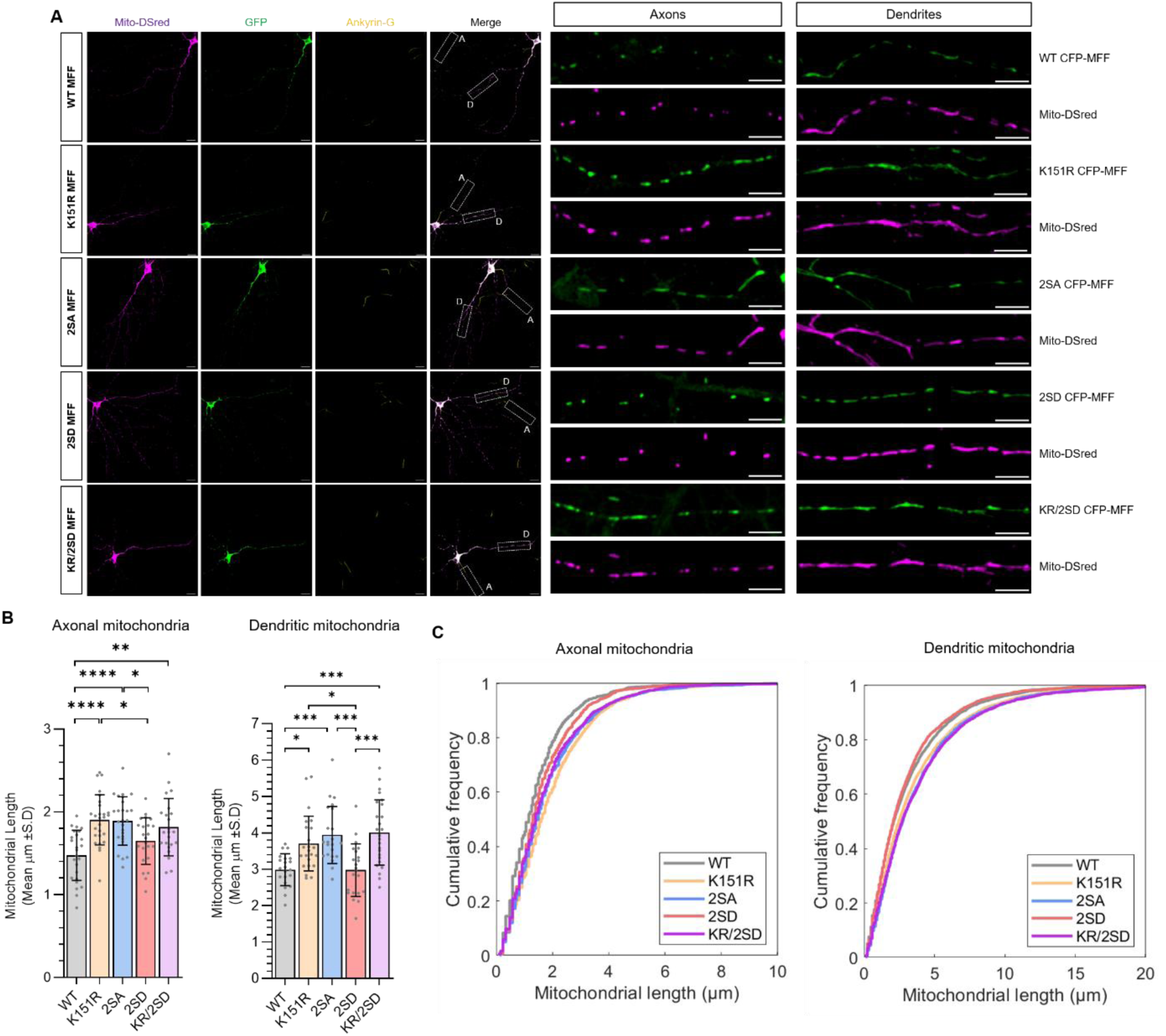
The MFF phosphorylation-SUMOylation axis regulates dendritic mitochondrial length under basal conditions. (**A**) Confocal images of primary hippocampal neurons transfected with mito-DSred and the indicated CFP-MFF constructs. Neurons were fixed and stained for GFP and ankyrin-G (to define the axonal initial segment), scale bar 20 µm. Dotted box in merged panel denotes the axonal (A) and dendritic (D) region in the enlargements, scale bar 10µm. (**B**) Quantification of mean mitochondrial lengths. Data gathered from four independent experiments (three for KR/2SD). One-Way ANOVA followed by Tukey’s post hoc test (axons), Kruskal Wallis test followed by Dunn’s multiple comparison test (dendrites) used to determine significance. p*<0.05, p**<0.005, p***<0.0005, p***<0.0001. (**C**) Cumulative frequency of all axonal and dendritic mitochondria.

As observed in Fig 2, neurons expressing K151R exhibit elongated axonal mitochondria compared to wild-type expressing cells (∼29.1% increase). Neurons expressing 2SA had comparable length to K151R cells, with an ∼28.2% increase compared to wild-type MFF. Neurons expressing 2SD had significantly shorter axonal mitochondria compared to K151R and 2SA cells. Importantly, neurons expressing KR/2SD exhibited an elongation phenotype comparable to the single mutation of K151R and the phospho-null 2SA, inducing an increase of ∼23.1% compared to wild-type MFF (Fig 3B).

Dendritic mitochondrial lengths followed a similar pattern to axonal mitochondria; wild-type and 2SD MFF expressing neurons exhibited similar mean mitochondrial lengths, whereas 2SA had significantly longer mitochondria compared to both wild-type and 2SD (increase of ∼28.4% and 32.1%, respectively). As in Fig 2, neurons expressing K151R had longer mitochondria compared to wild-type and 2SD (increase of ∼16.8% and ∼20.1%, respectively), and at comparable levels to 2SA. Strikingly, the KR/2SD mutant exhibited similar mitochondrial lengths to the 2SA and K151R mutant, and significantly longer mitochondria compared to wild-type and 2SD (increase of ∼28.3% and ∼31.9%, respectively, Fig 3B). The cumulative frequency graphs of axonal and dendritic mitochondria demonstrate a shift to longer mitochondria for K151R, 2SA and KR/2SD, which exhibit similar curves, compared to WT and 2SD (Fig 3C). These results demonstrate that SUMOylation of MFF initiated by phosphorylation is a key facilitator of fission, which regulates mitochondrial size in axons and dendrites under basal conditions.

### MFF SUMOylation is required for AICAR-induced mitochondrial fragmentation

Following the observation that MFF SUMOylation regulates mitochondrial fission under basal conditions, we performed morphological analysis on dendritic mitochondria of transfected neurons with GFP-MFF (wild-type or K151R) and mito-DsRed in neurons treated with AICAR (Fig 4A). Because our data indicate that AICAR does not impact axonal mitochondrial length (Fig 1B) we focused our analysis on dendritic mitochondria. As expected, neurons expressing wild-type MFF exhibited mitochondrial fragmentation following AICAR treatment (reduction of ∼17.9%) and a shift to shorter mitochondria (Fig 4C). However, cells expressing MFF-K151R were resistant to AICAR-induced fission, with no difference detected in average mitochondrial length between control and AICAR-treated cells (Fig 4B) or in cumulative frequency (Fig 4C). These results are consistent with the data shown in Figures 2 and 3, that MFF SUMOylation is required for AMPK-induced mitochondrial fission in dendrites. Together, these results demonstrate that MFF SUMOylation, following AMPK activation, drives dendritic mitochondrial fragmentation.

**Figure 4.**
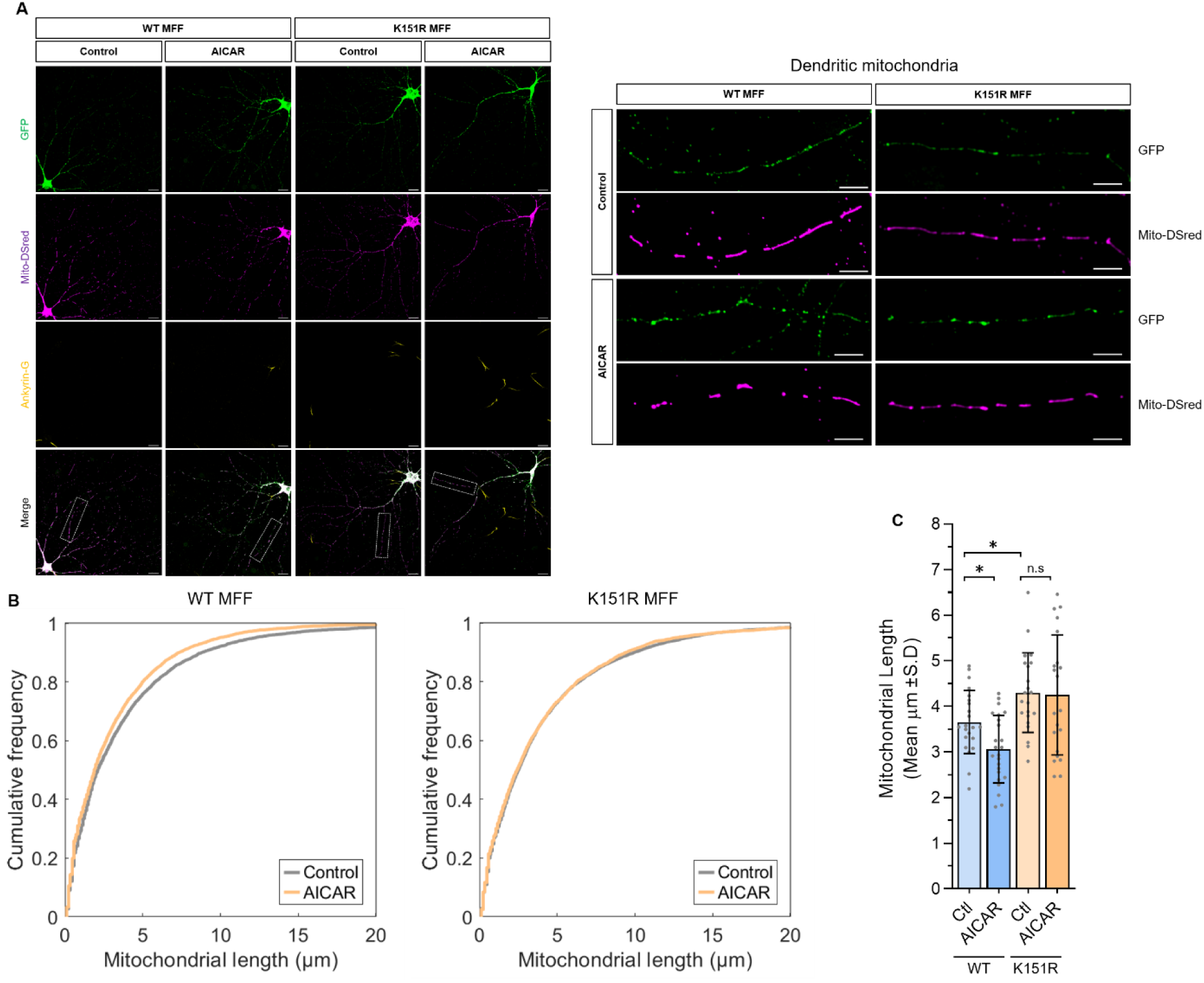
AICAR-induced mitochondrial fragmentation is MFF SUMOylation-dependent. (**A**) Confocal images of primary hippocampal neurons transfected with mito-DSred and either wild-type or K151R GFP-MFF constructs, and treated with 1mM AICAR for 3 days. Neurons were fixed and stained for ankyrin-G and GFP, scale bar 20 µm. Enlargements show highlighted region of dendrite, scale bar 10µm. (**B**) Cumulative frequency of all dendritic mitochondria. (**C**) Quantification of mean mitochondrial lengths. Data gathered from three independent experiments. Brown-Forsythe ANOVA test followed by Dunnett’s multiple comparison test used to determine significance. p*<0.05.

## Discussion

MFF is a key target of AMPK phosphorylation during bioenergetic stress (Toyama *et al*, 2016; Ducommun *et al*, 2015). We have previously demonstrated that AMPK phosphorylation of MFF leads to its subsequent SUMOylation at Lys^151^, which then promotes mitochondrial fragmentation in cell lines (Seager *et al*, 2024). In the current work, we focused on neurons to show that the AMPK-MFF phosphorylation-SUMOylation axis is a key pathway in both the basal regulation of axonal and dendritic mitochondria, in AMPK-mediated fragmentation of dendritic mitochondria. These results support and extend findings from a previous report indicating the importance of MFF phosphorylation in AMPK-induced dendritic mitochondrial fragmentation (Lee *et al*, 2022). Our data are consistent with a model in which activated AMPK phosphorylation of MFF at Ser^155^ and Ser^172^ enhances MFF SUMOylation to promote mitochondrial fission in neurons. This model is supported by the following observations: (1) expression of the SUMO-deficient MFF mutant K151R elongates mitochondria in axons and dendrites, (2) blocking SUMOylation in the MFF phospho-mimetic (K151R/2SD) phenocopies the elongation seen with K151R and 2SA (phospho-null) expression, (3) AMPK activation with AICAR fragments dendritic mitochondria in an MFF SUMO-dependent manner.

Consistent with previous reports (Chang *et al*, 2006; Lewis *et al*, 2018) we observe axonal mitochondria as sparce and punctate, whereas dendritic mitochondria are elongated and occupy more of the process. One core question is what differential signalling pathways underpin this compartment-specific mitochondrial morphology, since both MFF (Lewis *et al*, 2018) and AMPK (Watters *et al*, 2020) are present in axons and dendrites? There are conflicting reports in the literature about the effect of MFF knockdown on basal mitochondrial morphology. It has been reported that MFF knockdown does not affect dendritic mitochondrial length (Lee *et al*, 2022; Lewis *et al*, 2018) but specifically elongates axonal mitochondria (Lewis *et al*, 2018), whereas other groups have reported that MFF does regulate dendritic mitochondrial length (Hatsuda *et al*, 2023; Virga *et al*, 2024).

Lewis and colleagues speculate the axonal/dendritic difference they observe could be due to compartment-specific MFF activation, splice variants and post-translational modifications, altering interactions with DRP1 and hence mitochondrial size (Lewis *et al*, 2018). Indeed, different isoforms of MFF have different phosphorylation states and fission efficiencies (Hanada *et al*, 2024), highlighting that differential MFF activity may explain basal morphological differences. Intriguingly, it has been reported recently that within the same neuron, differences in the fusion and fission balance between dendrites proximal to the soma, and dendrites distal from the soma are due to differential AMPK activity (Virga *et al*, 2024). This proposal is consistent with the report that specific AMPK activation within either the axon or somatodendritic compartment regulates local mitochondria in a compartment-confined manner (Watters *et al*, 2020). Indeed, AMPK activity is synchronised to calcium spikes following neuronal activity in the somatodendritic compartment, not observed in the axon (Hatsuda *et al*, 2023). These observations highlight signalling variation between neuronal compartments, and AMPK activity is likely differentially regulated in dendrites and axons leading to diverse, compartment-specified mitochondrial morphologies. Differential signalling and SUMO pathway regulation between the compartments could be established resulting in different basal levels of MFF SUMOylation; in the axon MFF SUMOylation could be higher compared to the dendritic compartment, resulting in shorter mitochondria.

Our previous work identified that in MEF cells, under basal conditions, no difference in morphology was observed between WT and K151R MFF, (Seager *et al*, 2024). We propose this could be due to several reasons: (1) at a basal level, MFF may be more SUMOylated in neurons, therefore blocking MFF SUMOylation has a more apparent effect, (2) the other DRP1 receptors can compensate for the loss of MFF SUMOylation in MEF cells, and (3) the morphology of mitochondria in neurons is more ‘restricted’ to tubular structures within neurites, compared to the network-like architecture in fibroblasts, therefore observing changes in neurons is more apparent.

During neuronal activity in developing neurons, AMPK signalling is important for maintaining mitochondrial homeostasis. Preventing mitochondrial fission by silencing AMPK or MFF results in defective dendritic branch formation, impaired mitophagy and elongated mitochondria with reduced function (Hatsuda *et al*, 2023). AMPK signalling is also required during neuronal activity in mature neurons to drive mitochondrial fission in an MFF-dependent manner (Virga *et al*, 2024). Moreover, a recent study identified that shorter dendritic mitochondria have a reduced membrane potential, impaired Ca^2+^ uptake during neuronal activity and reduced dendritic spine density (Strucinska *et al*, 2025).

Correct dendritic mitochondrial morphology supports protein synthesis and spine plasticity (Rangaraju *et al*, 2019). During long-term potentiation (LTP), dendritic spines undergo structural changes and an increase in volume, accompanied by enhanced mitochondrial matrix Ca^2+^ levels and mitochondrial fission. Blocking mitochondrial fission prevents mitochondrial Ca^2+^ uptake and structural changes during LTP (Divakaruni *et al*, 2018). Although the mechanisms are not fully elucidated, the authors speculate that fission enhances mitochondrial Ca^2+^ handling capacity to increase mitochondrial function (Divakaruni *et al*, 2018). MFF-silencing significantly elongates mitochondria, enhances Ca^2+^ uptake and reduces neurotransmitter release (Lewis *et al*, 2018). Taken together, these studies emphasise the importance of mitochondrial morphology development and neuronal activity, and the roles of MFF SUMOylation in regulating mitochondrial size, and the impact on synaptic function. Could tuning the level of MFF SUMOylation exquisitely alter mitochondrial length, therefore alter Ca^2+^ buffering capacity, thus impacting neuronal function? Future studies should investigate the role of MFF SUMOylation in developing neurons and during neuronal activity.

Disrupted mitochondrial dynamics is a hallmark of many neurodegenerative disease (Sirois *et al*, 2026; Chen *et al*, 2023). The dynamic regulation of AMPK is important during neuronal activity to sustain energy levels and recruit mitochondria to synapses (Li *et al*, 2020; Marinangeli *et al*, 2018). Hyperactivated AMPK is observed in AD (Vingtdeux *et al*, 2011; Ma *et al*, 2014; Fang *et al*, 2019), and the toxic AD-associated protein Aβ over activates AMPK in primary neurons (Thornton *et al*, 2011; Mairet-Coello *et al*, 2013), which is required for Aβ-induced synaptic spine loss (Yang *et al*, 2022; Mairet-Coello *et al*, 2013; Lee *et al*, 2022; Domise *et al*, 2019). Aβ-induced synaptic plasticity defects are rescued upon inhibition of AMPK (Ma *et al*, 2014), revealing overactive AMPK signalling as a therapeutic target in AD. Here we used the AMPK activator AICAR to chronically activate AMPK in mature neurons for three days, resulting in MFF phosphorylation and fragmentation of dendritic mitochondria, with no change observed in the axonal compartment, agreeing with previous reports (Hatsuda *et al*, 2023).

Application of oligomeric Aβ causes mitochondrial fragmentation in the dendrites, by AMPK-mediated phosphorylation of MFF, leading to spine loss (Lee *et al*, 2022). Importantly, blocking mitochondrial fragmentation by either knockdown of MFF or expression of the phospho-null 2SA (as used in the current study) prevents mitochondrial fission and spine loss, indicating that AMPK-mediated phosphorylation of MFF is a prerequisite for Aβ-induced neuronal dysfunction (Lee *et al*, 2022). Our findings align with those of Lee and colleagues, and we demonstrate the importance of the AMPK-MFF phosphorylation axis in driving dendritic mitochondrial fission, and that MFF SUMOylation is key to this pathway. Indeed, blocking mitochondrial fragmentation in a number of neurodegenerative models is neuroprotective (Strucinska *et al*, 2025; Lee *et al*, 2022; Filichia *et al*, 2016; Bido *et al*, 2017; Qi *et al*, 2012; Grohm *et al*, 2012). Although not tested in the current work, future work should investigate synaptic spine loss under AICAR and Aβ application, and interrogate if blocking MFF SUMOylation is protective under these conditions.

Mitochondrial fission regulator 1-like protein (MTFR1L) is an OMM protein recently demonstrated to regulate mitochondrial morphology. Loss of MTFR1L causes mitochondrial elongation associated with an increase in the fusion protein OPA1 and more frequent fusion events, indicating MTFR1L is an anti-fusion protein. Similar to MFF, MTFR1L activity is modulated by AMPK. Direct AMPK activation or mitochondrial stress leads to AMPK phosphorylation of MTFR1L, which is necessary for mitochondria to undergo fragmentation by reducing OPA1 levels (Tilokani *et al*, 2022). In neurons, AMPK-mediated phosphorylation of MTFR1L and MFF are both required for fission of dendritic mitochondria (Virga *et al*, 2024). Interestingly, we observe no change in OPA1 levels in neurons treated with AICAR for three days (Fig 1F). Since MTFR1L would reduce OPA1 levels to drive fission, this observation argues, at least under the conditions and time point used here, that MFF is sufficient to promote fission. Since MTFR1L activity was not directly tested in the current work, we cannot rule out that MTFR1L activity contributed to the observed fragmentation, perhaps at an earlier time point. Future studies should investigate the regulation of both these proteins in AMPK-mediated mitochondrial remodelling.

In conclusion, we show AMPK-mediated MFF-phosphorylation-SUMOylation regulates basal mitochondrial morphology. Future work should investigate the relationship between MFF SUMOylation-mediated morphological changes and the impact on mitochondrial and synaptic function. We also identify that chronic AMPK activation fragments dendritic mitochondria in an MFF-SUMO-dependent manner. With the pathological relevance of excessive mitochondrial fragmentation and AMPK hyperactivation in AD, targeting MFF-SUMOylation offers a potential therapeutic avenue for neuroprotection.

## Materials and Methods

### Reagents and plasmids

Lipofectamine 2000 was from Invitrogen. Poly-L-lysine and poly-D-lysine (PLL, PDL) were from Sigma. AICAR (dissolved in cell culture-grade H2O) was from Tocris. For Western blotting, primary antibodies used were AMPK (ThermoFisher, cat # AHO1332), AMPK-P (Cell Signalling, cat # 2535), β actin (Sigma, cat # A5441), DRP1 (BD Biosciences, cat # 611113), MFF (Santa Cruz, SC-398731), MFF-P (Invitrogen, PA5-104614), OPA1 (Abcam, AB42364),

VDAC (Cell Signalling, D73D12). Secondary antibodies were horseradish peroxidase (HRP)– conjugated anti-mouse (raised in goat), anti-goat (raised in rabbit), anti-rabbit (raised in goat), obtained from Sigma-Aldrich, and used at a dilution of 1:10,000.

Primary antibodies used for imaging were anti-GFP (chicken, Abcam, cat # 13970, at 1:1000, to enhance CFP/GFP signal on expressed MFF constructs), and anti-ankyrin-G (mouse, Synaptic Systems, cat # 160103, 1:1000). Secondary antibodies for imaging were anti-chicken Cy2 (Jackson Labs, 1:1000) and anti-mouse Alexa Fluor 647 (Jackson Labs, 1:1000).

Constructs of MFF human isoform 1 were generated by subcloning the MFF human isoform 1 sequence from GST-MFF into the Bam HI/Hind III sites of pECFP-C1, and site directed mutagenesis used to generate the SUMO and phosphorylation mutations, as previously described (Seager *et al*, 2024). To generate the triple MFF-K151R/2SD mutant, we used site-directed mutagenesis of the lysine to arginine within the CFP-MFF 2SD construct using the same primers to generate the K151R, as before (Seager *et al*, 2024). MitoDS-red (pDsRed2-mito) was from Clontech.

### SDS-PAGE and Western blot analysis

Protein lysates were collected by washing once in ice cold PBS, lysing directly in 1x Laemmli buffer, and then boiling at 95°C for 10 min. SDS-PAGE was performed using standard Bio-Rad equipment and proteins resolved by 10% tris-glycine polyacrylamide gels, which were made in-house. Proteins were transferred to polyvinylidene difluoride membranes (Merck), blocked in PBS-T with either 5% milk or 4% bovine serum albumin for 1 hour at room temperature, and incubated with primary antibody (1 hour at room temperature or overnight at 4°C). Membranes were washed with PBS-T and incubated with secondary antibody at room temperature for 1hr. Membranes were washed in PBS-T and assayed for chemiluminescence using a Li-COR Odyssey Fc scanner. Densitometry analysis was performed using the Li-COR image analysis software.

### Neuronal culture and transfection

Primary hippocampal and cortical neurons were dissociated and cultured from Han Wistar rats at embryonic day 17, as previously described (Nair *et al*, 2021). For biochemistry, 500,000 cortical neurons were cultured in 6-well, or 125,000 neurons for 24-well plastic plates, precoated with PLL. For imaging, 100,000-150,000 hippocampal neurons were plated onto 22 mm PDL coated glass coverslips. Cells were plated into plating media (Neurobasal® (Gibco) supplemented with 5% horse serum, 2% B27, 1% glutamax, and 1% penicillin-streptomycin) maintained at 37°C/5% CO2. The following day, the media was fully exchanged for serum free media (feeding media: Neurobasal® medium supplemented with 2% B27, 1% glutamax and 1% penicillin-streptomycin) and maintained at 37°C/5% CO2. Cells were supplemented with 0.5ml warmed feeding media on DIV 7 and DIV14.

Neurons were transfected with 1 µg of each plasmid DNA and 1.5 µL lipofectamine 2000 per 1 µg DNA according to manufactures instructions, with some modifications. Coverslips were carefully removed from culture dishes, briefly washed in warmed plain neurobasal media, and placed in transfection media (feeding media lacking antibiotics). DNA and lipofectamine complexes were prepared in plain neurobasal media, incubated at room temperature for 20 minutes, and added dropwise to the coverslips, and returned to the incubator for 45-60 minutes. Coverslips were carefully removed, washed in plain media and returned to the original culture dishes for 2-3 days. For figures 2-3, transfections occurred at DIV 12-15, and cells transfected for 2-3 days until fixation. For figures 1 and 4 (AICAR treatments) transfections occurred on DIV 15, cells incubated for 3 days.

### Immunocytochemistry

Transfected neurons were washed briefly in warm PBS and fixed in 4% PFA/PBS for 15 mins at room temperature, washed three times in PBS and then permeabilized with 0.1% Triton X-100/PBS for 3-4 min, and then washed in PBS. To quench unreacted formaldehyde, cells were incubated in 100 mM glycine/PBS for 3-4 min, and then washed once in PBS. Cells were incubated in 3% BSA/PBS for 20 min at room temperature to block non-specific binding. Primary antibody (GFP, 1:1000, Ankyrin G 1:1000) was prepared in 3% BSA/PBS, and coverslips were incubated with primary antibody for 60 min at room temperature. Coverslips were washed three times with PBS and then incubated with secondary antibody (prepared in 3% BSA/PBS at 1:1000) for 45 min. Coverslips were washed four times in PBS and mounted on glass microscope slides using Fluoromount-G containing DAPI.

### Mitochondrial morphology analysis

Confocal images were taken using a Leica SP5 inverted microscope with a 63x lens. 0.5µm z-slices were taken of the cell body, axonal and dendritic regions. Images were analysed using ImageJ software. Firstly, ROIs for the axon and dendrites were generated using the freehand tool and saved (the ankyrin G stain was used to differentiate between the axon and dendritic processes). Images were max projected, ‘tubeness’ analysis applied (sigma: 0.2045), converted to 8-bit, and a threshold (Li method) applied. The image was then skeletonised and the ROI’s applied to define only the axons and dendrites. Skeletonised images were manually checked against the original confocal image to ensure accurate representation. Using the ‘analyse skeleton’ function, branch length data was extracted as mitochondrial length. Any mitochondria appearing as a single pixel is not considered a branch using the analyse skeleton function, and appears in the branch length with zero length. These were manually included in the analysis and given a length of 0.12um (half of the mitochondrial length which appear as 2 pixels).

The cumulative distribution functions were generated using a custom script written in MATLAB (version R2024a). For each genotype and condition all mitochondrial lengths were pooled together and their cumulative distribution function was plotted.

### Mitochondrial ATP measurements

ATP levels were measured using the Mitochondrial ToxGlo™ Assay from Promega, following manufacturers protocols. Briefly, 200uL of media from each well (cells cultured in 12-well plates) was removed and added to an equal volume of 2X ATP detection reagent and carefully mixed. The remaining media was aspirated, and 350µL/well of ATP detection mixture added back to the cells (ensuring sufficient media-ATP detection mixture was reserved for background measurement), and incubated at room temperature on an orbital shaker for 5 minutes. 100µL of ATP reagent mixture was transferred to 96-well opaque walled plates, each well plated in triplicate, and measured using the Mitochondrial ToxGlo ATP Assay on a Promega GloMax® Explorer microplate reader. Triplicate wells were averaged and background subtracted.

### Statistical analysis

Statistical analysis was performed using GraphPad Prism software version 9. Only experiments repeated at least 3 independent times were considered for statistical analysis. For quantification of Western blots, all values are presented as means ±SD, expressed as a percentage of control. One-sample *t* test was used to determine significance between conditions and control. For analysis of imaging, data were tested for normality distribution using the D’Agostino and Pearson test. If this test was passed, a t sample t test, or one-way ANOVA followed by Tukey’s post hoc test was used to determine significance. If failed, a Mann-Whitney U or Kruskal-Wallis test followed by Dunn’s post hoc test was used. For unevenly distributed data, the Brown-Forsythe ANOVA test followed by Dunnett’s T3 post hoc test was used. A *p* value of <0.05 was considered significant. *P* values, independent repeats and statistical approach are described in the figure legends.

## Author contributions

Conceptualisation: R.S., K.A.W

Methodology: R.S.

Experimentation: R.S.

Data analysis: R.S. and D.F

Supervision: K.A.W. and J.M.H.

Writing: original draft: R.S.

Editing: All authors contributed to editing the manuscript.

## Competing interests

The authors declare that they have no competing interests.

## Funding

This work was supported by the BBSRC (BB/R00787X/1), Wellcome Trust (105384/Z/14/A), Leverhulme Trust (RPG-2019-191), and the Academy of Medical Sciences Springboard (SBF0010\1086).

---

**Supplementary Figure 1.**
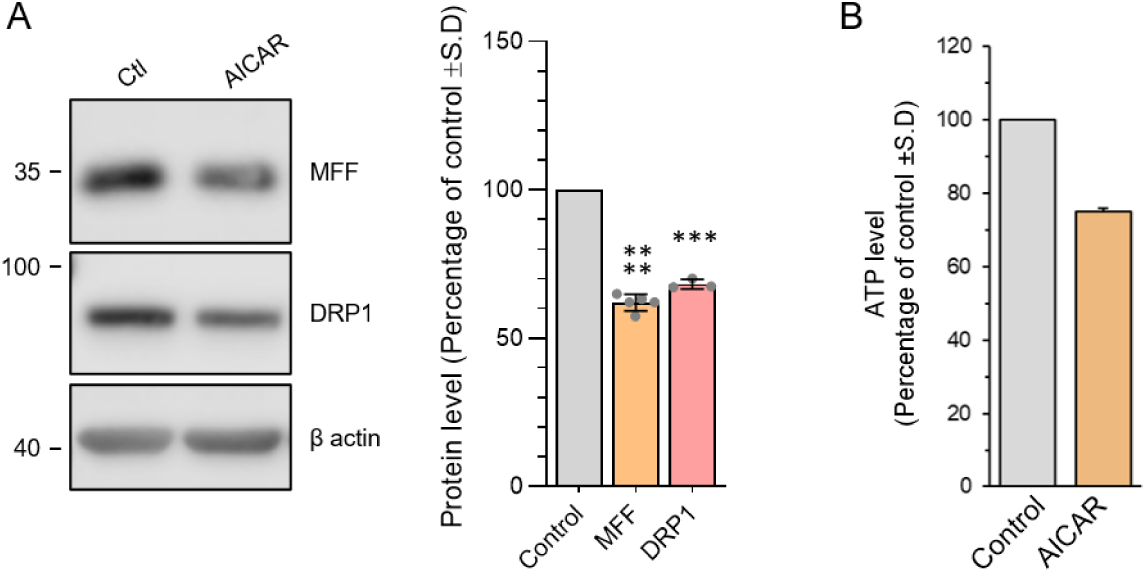
(**A**) AICAR-treated cortical neurons were probed for total levels of MFF and DRP1. Quantification shows data normalised to β-actin and expressed as percentage of control. Data generated from 3-5 independent experiments. One sample t test, p***<0.005, p****<0.001. (**B**) ATP assay of neurons grown in low glucose. Cortical neurons were cultured in 5 mM glucose from DIV1. On DIV18 cells were treated with 1 mM AICAR for three days and ATP assay performed. Data generated from two independent experiments.

## Notes

### Competing Interest Statement

The authors have declared no competing interest.

## References

Anand R, Wai T, Baker MJ, Kladt N, Schauss AC, Rugarli E & Langer T (2014) The i-AAA protease YME1L and OMA1 cleave OPA1 to balance mitochondrial fusion and fission. Journal of Cell Biology 204: 919–929

Bapat O, Purimetla T, Kruessel S, Shah M, Fan R, Thum C, Rupprecht F, Langer JD & Rangaraju V (2024) VAP spatially stabilizes dendritic mitochondria to locally support synaptic plasticity. Nat Commun 15

Bido S, Soria FN, Fan RZ, Bezard E & Tieu K (2017) Mitochondrial division inhibitor-1 is neuroprotective in the A53T-α-synuclein rat model of Parkinson’s disease. Sci Rep 7: 7495

Cagalinec M, Safiulina D, Liiv M, Liiv J, Choubey V, Wareski P, Veksler V & Kaasik A (2013) Principles of the mitochondrial fusion and fission cycle in neurons. J Cell Sci 126: 2187–2197

Chang DTW, Honick AS & Reynolds IJ (2006) Mitochondrial Trafficking to Synapses in Cultured Primary Cortical Neurons. Journal of Neuroscience 26: 7035–7045

Chen W, Zhao H & Li Y (2023) Mitochondrial dynamics in health and disease: mechanisms and potential targets. Signal Transduct Target Ther 8: 333

Datta S & Jaiswal M (2021) Mitochondrial calcium at the synapse. Mitochondrion 59: 135– 153

Devine MJ & Kittler JT (2018) Mitochondria at the neuronal presynapse in health and disease. Nat Rev Neurosci 19: 63–80

Divakaruni SS, Van Dyke AM, Chandra R, LeGates TA, Contreras M, Dharmasri PA, Higgs HN, Lobo MK, Thompson SM & Blanpied TA (2018) Long-Term Potentiation Requires a Rapid Burst of Dendritic Mitochondrial Fission during Induction. Neuron 100: 860–875

Domise M, Sauvé F, Didier S, Caillerez R, Bégard S, Carrier S, Colin M, Marinangeli C, Buée L & Vingtdeux V (2019) Neuronal AMP-activated protein kinase hyper-activation induces synaptic loss by an autophagy-mediated process. Cell Death Dis 10: 221

Ducommun S, Deak M, Sumpton D, Ford RJ, Núñez Galindo A, Kussmann M, Viollet B, Steinberg GR, Foretz M, Dayon L, et al (2015) Motif affinity and mass spectrometry proteomic approach for the discovery of cellular AMPK targets: Identification of mitochondrial fission factor as a new AMPK substrate. Cell Signal 27: 978–988

Fang EF, Hou Y, Palikaras K, Adriaanse BA, Kerr JS, Yang B, Lautrup S, Hasan-Olive MM, Caponio D, Dan X, et al (2019) Mitophagy inhibits amyloid-β and tau pathology and reverses cognitive deficits in models of Alzheimer’s disease. Nat Neurosci 22: 401–412

Filichia E, Hoffer B, Qi X & Luo Y (2016) Inhibition of Drp1 mitochondrial translocation provides neural protection in dopaminergic system in a Parkinson’s disease model induced by MPTP. Sci Rep 6: 32656

Grohm J, Kim S-W, Mamrak U, Tobaben S, Cassidy-Stone A, Nunnari J, Plesnila N & Culmsee C (2012) Inhibition of Drp1 provides neuroprotection in vitro and in vivo. Cell Death Differ 19: 1446–1458

Hanada Y, Maeda R, Ishihara T, Nakahashi M, Matsushima Y, Ogasawara E, Oka T & Ishihara N (2024) Alternative splicing of Mff regulates AMPK-mediated phosphorylation, mitochondrial fission and antiviral response. Pharmacol Res 209: 107414

Hatsuda A, Kuris J, Fujishima K, Kawaguchi A, Ohno N & Kengaku M (2023) Calcium signals tune AMPK activity and mitochondrial homeostasis in dendrites of developing neurons. Development 150: dev201930.

Herzig S & Shaw RJ (2018) AMPK: guardian of metabolism and mitochondrial homeostasis. Nat Rev Mol Cell Biol 19: 121–135

Lee A, Kondapalli C, Virga DM, Lewis TL, Koo SY, Ashok A, Mairet-Coello G, Herzig S, Foretz M, Viollet B, et al (2022) Aβ42 oligomers trigger synaptic loss through CAMKK2-AMPK-dependent effectors coordinating mitochondrial fission and mitophagy. Nat Commun 13: 4444

Lewis TL, Kwon SK, Lee A, Shaw R & Polleux F (2018) MFF-dependent mitochondrial fission regulates presynaptic release and axon branching by limiting axonal mitochondria size. Nat Commun 9: 5008

Li S, Xiong G-J, Huang N & Sheng Z-H (2020) The cross-talk of energy sensing and mitochondrial anchoring sustains synaptic efficacy by maintaining presynaptic metabolism. Nat Metab 2: 1077–1095

Li Z, Okamoto K-I, Hayashi Y & Sheng M (2004) The Importance of Dendritic Mitochondria in the Morphogenesis and Plasticity of Spines and Synapses. Cell 119: 873–887

Losón OC, Song Z, Chen H & Chan DC (2013) Fis1, Mff, MiD49, and MiD51 mediate Drp1 recruitment in mitochondrial fission. Mol Biol Cell 24: 659–667

Ma T, Chen Y, Vingtdeux V, Zhao H, Viollet B, Marambaud P & Klann E (2014) Inhibition of AMP-Activated Protein Kinase Signaling Alleviates Impairments in Hippocampal Synaptic Plasticity Induced by Amyloid. Journal of Neuroscience 34: 12230–12238

Mairet-Coello G, Courchet J, Pieraut S, Courchet V, Maximov A & Polleux F (2013) The CAMKK2-AMPK Kinase Pathway Mediates the Synaptotoxic Effects of Aβ Oligomers through Tau Phosphorylation. Neuron 78: 94–108

Marinangeli C, Didier S, Ahmed T, Caillerez R, Domise M, Laloux C, Bégard S, Carrier S, Colin M, Marchetti P, et al (2018) AMP-Activated Protein Kinase Is Essential for the Maintenance of Energy Levels during Synaptic Activation. iScience 9: 1–13

Nair JD, Henley JM & Wilkinson KA (2021) Surface biotinylation of primary neurons to monitor changes in AMPA receptor surface expression in response to kainate receptor stimulation. STAR Protoc 2: 100992

Obashi K & Okabe S (2013) Regulation of mitochondrial dynamics and distribution by synapse position and neuronal activity in the axon. European Journal of Neuroscience 38: 2350–2363

Otera H, Miyata N, Kuge O & Mihara K (2016) Drp1-dependent mitochondrial fission via MiD49/51 is essential for apoptotic cristae remodeling. J Cell Biol 212: 531–544

Qi X, Qvit N, Su Y-C & Mochly-Rosen D (2012) Novel Drp1 inhibitor diminishes aberrant mitochondrial fission and neurotoxicity. J Cell Sci 126: 789–802

Rangaraju V, Lauterbach M & Schuman EM (2019) Spatially Stable Mitochondrial Compartments Fuel Local Translation during Plasticity. Cell 176: 73–84

Seager R, Lee L, Henley JM & Wilkinson KA (2020) Mechanisms and roles of mitochondrial localisation and dynamics in neuronal function. Neuronal Signal 4 doi:10.1042/NS20200008 [PREPRINT]

Seager R, Ramesh NS, Cross S, Guo C, Wilkinson KA & Henley JM (2024) SUMOylation of MFF coordinates fission complexes to promote stress-induced mitochondrial fragmentation. Sci Adv 10

Sirois CL, Lee J, Chambers AL & Zhao X (2026) Mitochondrial dynamics in neurodevelopment and neurodevelopmental disorders. Nat Rev Neurosci 27: 307–326

Strucinska K, Kneis P, Pennington T, Cizio K, Szybowska P, Morgan A, Weertman J & Lewis TL (2025) A role for Fis1 in dendritic development. Sci Rep 16: 3594

Sun T, Qiao H, Pan P-Y, Chen Y & Sheng Z-H (2013) Motile Axonal Mitochondria Contribute to the Variability of Presynaptic Strength. Cell Rep 4: 413–419

Thornton C, Bright NJ, Sastre M, Muckett PJ & Carling D (2011) AMP-activated protein kinase (AMPK) is a tau kinase, activated in response to amyloid β-peptide exposure. Biochemical Journal 434: 503–512

Tilokani L, Russell FM, Hamilton S, Virga DM, Segawa M, Paupe V, Gruszczyk A V., Protasoni M, Tabara L-C, Johnson M, et al (2022) AMPK-dependent phosphorylation of MTFR1L regulates mitochondrial morphology. Sci Adv 8

Tong BC-K, Gubinelli F, Burbulla LF & Harbauer AB (2026) Mitochondrial specialization and signaling shape neuronal function. Trends Neurosci 49: 141–154

Toyama EQ, Herzig S, Courchet J, Lewis TL, Loson OC, Hellberg K, Young NP, Chen H, Polleux F, Chan DC, et al (2016) AMP-activated protein kinase mediates mitochondrial fission in response to energy stress. Science (1979) 351: 275–281

Vaccaro V, Devine MJ, Higgs NF & Kittler JT (2017) Miro1-dependent mitochondrial positioning drives the rescaling of presynaptic Ca2+ signals during homeostatic plasticity. EMBO Rep 18: 231–240

Verstreken P, Ly C V., Venken KJT, Koh T-W, Zhou Y & Bellen HJ (2005) Synaptic Mitochondria Are Critical for Mobilization of Reserve Pool Vesicles at Drosophila Neuromuscular Junctions. Neuron 47: 365–378

Vingtdeux V, Davies P, Dickson DW & Marambaud P (2011) AMPK is abnormally activated in tangle-and pre-tangle-bearing neurons in Alzheimer’s disease and other tauopathies. Acta Neuropathol 121: 337–349

Virga DM, Hamilton S, Osei B, Abigail M, Parker K, Zamponi E, Park JN, Hewitt LV, Zhang D, Gonzalez KC, et al (2024) Activity-dependent compartmentalization of dendritic mitochondria morphology through local regulation of fusion-fission balance in neurons in vivo. 8: 2142

Walters GC & Usachev YM (2023) Mitochondrial calcium cycling in neuronal function and neurodegeneration. Front Cell Dev Biol 11

Wang L, Guo L, Lu L, Sun H, Shao M, Beck SJ, Li L, Ramachandran J, Du Y & Du H (2016) Synaptosomal Mitochondrial Dysfunction in 5xFAD Mouse Model of Alzheimer’s Disease. PLoS One 11: e0150441

Wang X, Su B, Siedlak SL, Moreira PI, Fujioka H, Wang Y, Casadesus G & Zhu X (2008) Amyloid-β overproduction causes abnormal mitochondrial dynamics via differential modulation of mitochondrial fission/fusion proteins. Proceedings of the National Academy of Sciences 105: 19318–19323

Watters O, Connolly NMC, König H-G, Düssmann H & Prehn JHM (2020) AMPK Preferentially Depresses Retrograde Transport of Axonal Mitochondria during Localized Nutrient Deprivation. The Journal of Neuroscience 40: 4798–4812

Yang AJT, Mohammad A, Tsiani E, Necakov A & MacPherson REK (2022) Chronic AMPK Activation Reduces the Expression and Alters Distribution of Synaptic Proteins in Neuronal SH-SY5Y Cells. Cells 11: 2354

